# Validation of an LC-MS/MS Assay for the Simultaneous Determination of Bictegravir, Doravirine, and Raltegravir in Human Plasma

**DOI:** 10.1101/2020.05.21.104125

**Authors:** Amanda P. Schauer, Craig Sykes, Mackenzie L Cottrell, Angela DM Kashuba

**Affiliations:** Division of Pharmacotherapy and Experimental Therapeutics, University of North Carolina at Chapel Hill, Chapel Hill, North Carolina 27599, USA

## Abstract

Bictegravir (BIC), an integrase inhibitor, and doravirine (DOR), a non-nucleoside reverse transcriptase inhibitor, were recently approved by the US FDA for HIV treatment and are recommended first line treatment options. Because certain clinical scenarios warrant using them in combination, we developed a fully validated LC-MS/MS method for simultaneous measurement of BIC and DOR, along with a legacy integrase inhibitor, raltegravir (RAL), in human plasma over a clinically relevant 1000-fold range for each analyte. These analytes were extracted from the plasma by protein precipitation with their stable, isotopically labeled internal standards (BIC-d_5_, ^13^C_6_-DOR, and RAL-d_6_). Following extraction, samples were analyzed by reverse phase chromatography on a Waters Atlantis T3 C18 (50×2.1mm, 3um particle size) column with subsequent detection by electrospray ionization in positive ion mode on an AB Sciex API-5000 triple quadrupole mass spectrometer. The assay was linear (R^2^>0.994) over the selected calibration ranges (20.0-20,000ng/mL (BIC), 3.00-3,000ng/mL (DOR), and 10.0-10,000 (RAL)). The assay was accurate (inter-assay %Bias ≤±8.5) and precise (inter-assay %CV ≤11.4). This method was validated according to FDA guidance for industry and can be used to assess the pharmacokinetics of two newly approved antiretrovirals, or to support therapeutic drug monitoring for modern antiretroviral therapy.

## 1. Introduction

Since the introduction of highly active antiretroviral therapy (HAART) for HIV treatment in 1995, there remains the approach of using 2 or 3 mechanistically distinct antiretrovirals (ARVs) for durably suppressing HIV replication [1]. Therefore, it is important to have the ability to quantify multiple ARVs in patient specimens (such as plasma) for therapeutic drug monitoring or pharmacologic assessment of ARV combinations. ARVs encompass a wide range of chemical properties from extremely hydrophilic (i.e. nucleoside reverse transcriptase inhibitors; NRTIs) to extremely hydrophobic (i.e. protease inhibitors) making developing multiplex HPLC-MS/MS assays challenging. We sought to build on existing HPLC-MS/MS methods for 2 newly FDA approved first line antiretrovirals, bictegravir (an integrase strand transfer inhibitor; INSTI) [2,3] and doravirine (a nonnucleoside reverse transcriptase inhibitor; NNRTI) [3,4] by combining them into a single assay along with a legacy INSTI, raltegravir. This method needed to encompass a clinically relevant concentration range for each analyte.

## 2. Materials and Methods

### 2.1 Materials

BIC (purity, 98.27%) was purchased from MedChem Express (Monmouth Junction, NJ, USA), while its isotopically-labelled internal standard Bictegravir-d_5_ (BIC-d_5_) (purity, 97.5%) was purchased from AptoChem (Montreal, QC, Canada). Doravirine (purity, 99.5%) and its isotopically-labelled internal standard ^13^C_6_-Doravirine (^13^C_6_-DOR) (purity 95.1%) as well as the isotopically-labelled internal standard for Raltegravir (Raltegravir-d_6_, RAL-d_6_, purity 96.9%) were purchased from AlsaChim (Illkirch-Graddenstaden, France). Raltegravir (RAL) (purity 90.3%, including the potassium salt correction) was purchased with potassium salt from Toronto Research Chemicals (North York, ON, Canada). See Figure 1 for these chemical structures. Dimethyl sulfoxide (DMSO) (certified ACS), methanol (MeOH), acetonitrile (ACN) (both HPLC grade), and formic acid (certified ACS, 88%) were purchased from Fisher Scientific (Fair Lawn, NJ). Water used was purified by a Hydro Picosystem® UV Plus (Durham, NC). Human blank EDTA plasma used as a control throughout the method and whole blood used for a validation test were purchased from BioIVT (Westbury, NY).

**Figure 1:**
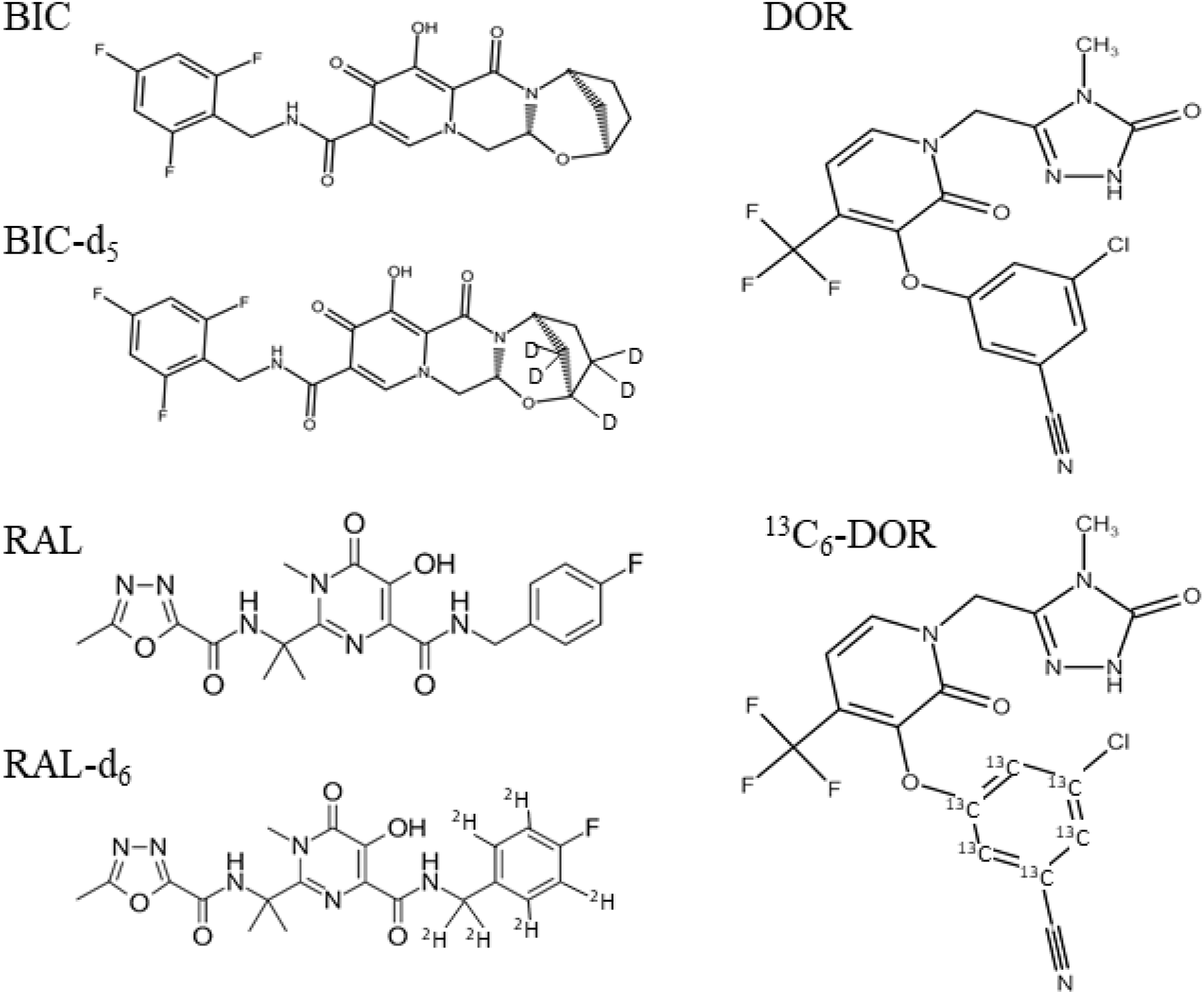
Chemical Structure of BIC, BIC-d_5,_ DOR, ^13^C_6_-DOR, RAL, and RAL-d_6_

### 2.2. LC-MS/MS Instrumentation

A Shimadzu Prominence HPLC system including pumps (LC-20AD), degasser (DGU-20A), and controller (CBM-20A) were supplied from Shimadzu (Columbia, MD, USA). A Waters Atlantis T3 C18 (50×2.1mm, 3um particle size) analytical column was used under reverse-phase conditions with 0.1% formic acid in water (mobile phase A) and 0.1% formic acid in acetonitrile (mobile phase B) at a flow rate of 0.500mL/min. The column heater (CTO-20A) was set at 40°C, and the autosampler (SIL-20AC/HT) was maintained at 10 °C. The injection volume was 2uL for the assay. The LC gradient held at 40% B for 0.25 minutes and increased to 90% B at 1.50 minutes and held for 30 seconds. At 2.10 minutes the gradient returned to 40% B until a final run time of 3.00 minutes. The retention time for BIC and BIC-d_5_ was 1.12 minutes, for DOR and ^13^C_6_-DOR was 1.28 minutes, and RAL and RAL-d_6_ was 1.04 minutes.

An API-5000 triple quadruple mass spectrometer (SCIEX, Foster City, CA, USA) operated in positive ion TurboIonspray mode was used to acquire data for this method. The source temperature was 500 °C and the ion spray voltage was 4500V. The declustering potential (DP)/collision energy (CE) for analyte/internal standard were as follows: BIC, BIC-d_5_ (216/59) DOR, ^13^C_6_-DOR (130/33), and RAL, RAL-d_6_ (131/25). Multiple reaction monitoring (MRM) was used to detect the analyte [precursor/product] transitions (m/z) as follows: BIC [450.2/261], BIC-d_5_ [455.2/266], DOR [428.0/112.0], ^13^C_6_-DOR [432.0/112.0], RAL [446.1/362.0], and RAL-d_6_ [451.1/367.0]. Natural isotopes of DOR (^37^Cl) and RAL (^13^C) were used to lower the ion intensity to achieve similar peak heights for all compounds at a given concentration.

Linear regression of concentration (x) versus peak area ratio of compound to internal standard (y) using a 1/(x^2^) weighting was used with Sciex Analyst software (version 1.6.2).

### 2.3. Validation

This analytical method was validated to meet the acceptance criteria of the US Food and Drug Administration guidelines [5]. The validation procedure consisted of three analytical runs to evaluate assay linearity, precision and accuracy, with additional runs to determine specificity, dilutions, stability, recovery, and matrix effects.

### 2.4. Preparation of Calibration Standards and Quality Control Samples

BIC, DOR, and RAL were weighed in duplicate and prepared to 1mg/mL in DMSO (BIC, DOR) or purified water (RAL). These 1mg/mL stocks were then diluted in 50:50 methanol:water to make a set of calibration standard working solutions at 200, 400, 1,000, 4,000, 10,000, 40,000, 170,000, 200,000 ng/mL (BIC), 30.0, 60.0, 150, 600, 1,500, 6,000, 25,500, 30,000 ng/mL (DOR), and 100, 200, 500, 2,000, 5,000, 20,000, 85,000, 100,000 ng/mL (RAL). Quality control (QC) working solution sets were also prepared in the same diluent with concentrations at 200, 600, 8,000, 160,000, 600,000 (dilution) ng/mL (BIC), 30.0, 90.0, 1,200, 24,000, 90,000 (dilution) ng/mL (DOR), and 100, 300, 4,000, 80,000, 300,000 (dilution) ng/mL (RAL). The stocks as well as the working solutions were stored at −80 °C.

Calibration standards and QCs were prepared fresh daily by spiking the appropriate working solution into human plasma. The resulting concentrations for the calibration standards were 20.0, 40.0, 100, 400, 1,000, 4,000, 17,000, 20,000 ng/mL (BIC), 3.00, 6.00, 15.0, 60.0, 150, 600, 2,550, 3,000 ng/mL (DOR), and 10.0, 20.0, 50.0, 200, 500, 2,000, 8,500, 10,000 ng/mL (RAL). QCs were prepared at the LLOQ, low QC, mid QC, high QC, and dilution QC concentrations as follows: 20.0, 60.0, 800, 16,000, 60,000 ng/mL (BIC), 3.00, 9.00, 120, 2,400, 9,000 ng/mL (DOR), and 10.0, 30.0, 400, 8,000, 30,000 ng/mL (RAL).

The internal standards, BIC-d_5_ and ^13^C_6_-DOR were prepared to 1 mg/mL in DMSO, while RAL-d_6_ was prepared to 0.500 mg/mL in purified water and all were stored at −80 °C. These solutions were further diluted with methanol into a single working solution with concentrations of 150, 15.0, and 75.0 ng/mL (BIC-d_5_, ^13^C_6_-DOR, RAL-d_6_) and were stored at 4 °C.

### 2.5. Sample Extraction

Human plasma (30μL) samples were extracted by protein precipitation by mixing with 270μL of methanol containing the stable, isotopically labeled internal standards (BIC-d_5_, ^13^C_6_-DOR, RAL-d_6_). The samples were then vortexed, centrifuged, and transferred into a 96-well plate for LC-MS/MS analysis.

## 3. Results and Discussion

### 3.1. Selectivity

Selectivity was evaluated in six unique lots of human plasma. No interfering peaks were detected at the retention time of the analytes or internal standard in any of the plasma lots evaluated. Representative chromatograms for matrix blanks, blanks with internal standard, and a LLOQ standard are shown in Figures 2–4.

**Figure 2:**
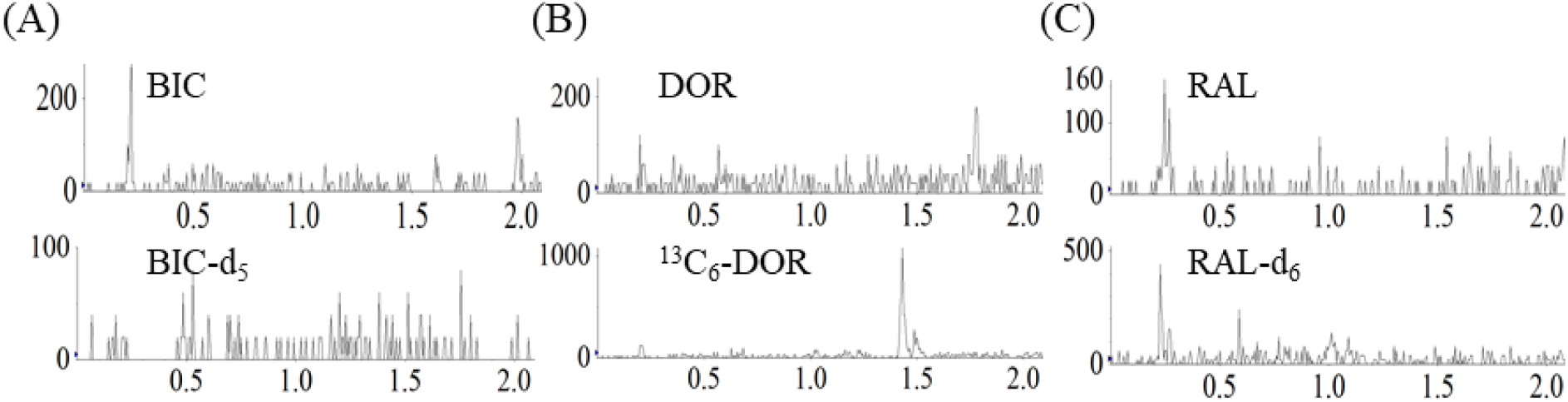
Example chromatograms from a blank for BIC, BIC-d_5_ (Figure 2A), DOR, ^13^C_6_-DOR (Figure 2B), and RAL, RAL-d_6_ (Figure 2C) in human plasma

**Figure 3:**
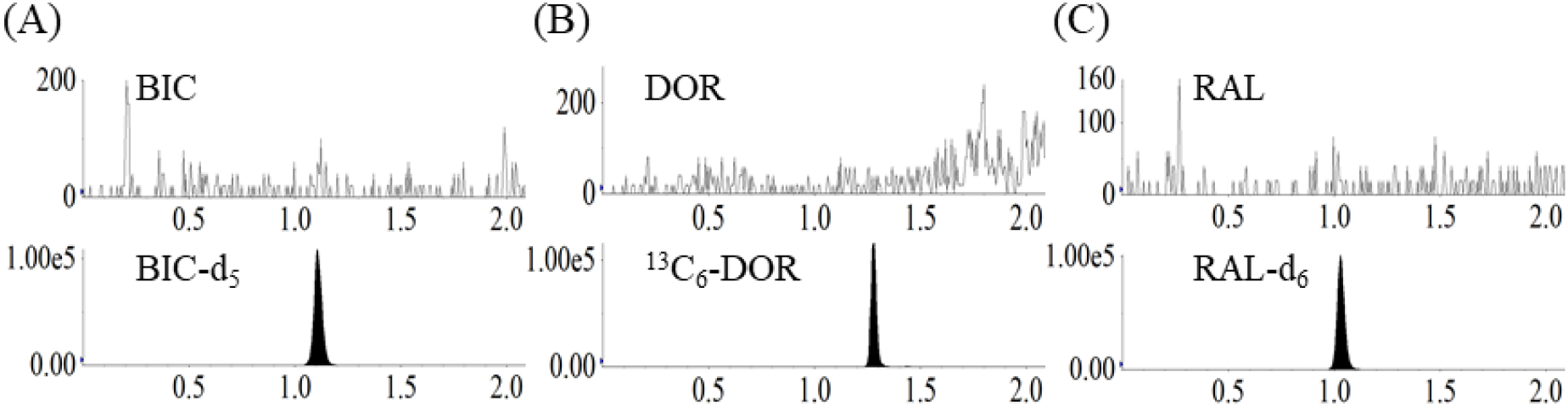
Example chromatograms from a blank with internal standard for BIC, BIC-d_5_ (Figure 3A), DOR, ^13^C_6_-DOR (Figure 3B), and RAL, RAL-d_6_ (Figure 3C) in human plasma

**Figure 4:**
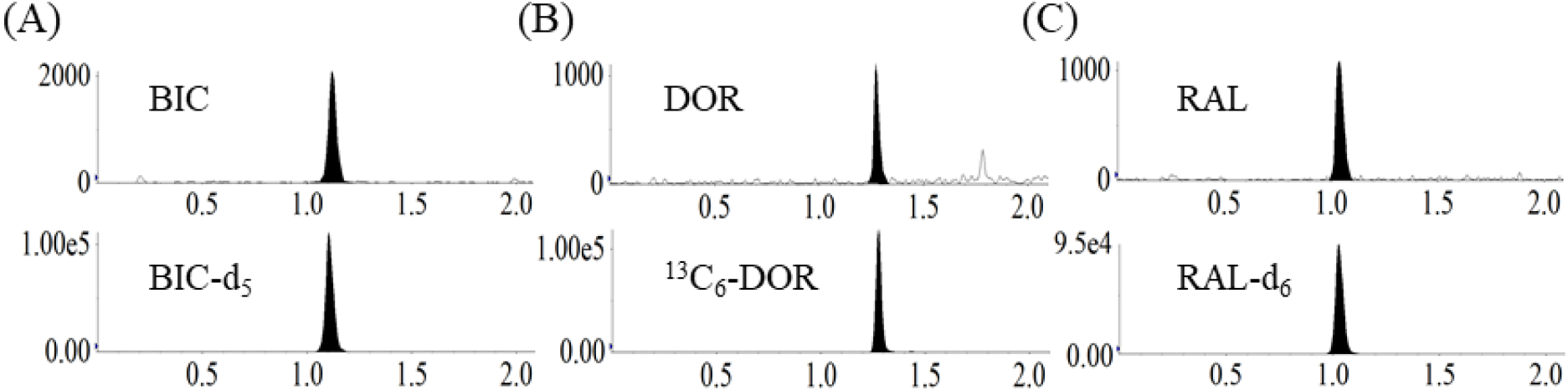
Example chromatograms from a LLOQ standard for BIC, BIC-d_5_ (Figure 4A), DOR, ^13^C_6_-DOR (Figure 4B), and RAL, RAL-d_6_ (Figure 4C) in human plasma

### 3.2. Linearity

Three accuracy and precision runs were extracted during this method validation. All calibration curves employed linear regression with 1/x^2^ weighting and correlation coefficients exceeded 0.99. Calibration standards were included at the beginning and the end of each analytical run to bracket all injected QCs. Peak area ratios of the calibration standard to internal standard were used to construct the calibration curves. All runs passed according to our predefined acceptance criteria for calibration standards of accuracy ≤ ±15% (≤ ±20% at the LLOQ).

### 3.3. Accuracy and Precision

Accuracy and precision were evaluated by analysis of six replicates of human plasma QC samples prepared at the LLOQ (20.0 ng/mL (BIC), 3.00 ng/mL (DOR) and 10.0 ng/mL (RAL)) and at three additional concentrations (60.0, 800, 16,000 ng/mL (BIC), 9.00, 120, 2,400 ng/mL (DOR), and 30.0, 500, 8,000 ng/mL (RAL)) over four runs. Inter-assay statistics were determined from replicate analysis of QC samples (n=6) in each of the 4 separate analytical runs (n=24). Precision was calculated as coefficient of variation (%CV) while accuracy was calculated as % Bias from the nominal concentration. Both met the predefined acceptance criteria of ±15% (±20% at the LLOQ). Inter-assay accuracy and precision results are shown in Table 1.

**Table 1:** Precision and Accuracy Results for BIC, DOR, and RAL in Human Plasma Quality Control Samples

| Analyte | Nominal conc.<br>(ng/mL) | Inter-assay Accuracy<br>(%Bias) | Inter-assay Precision<br>(%CV) |
| --- | --- | --- | --- |
| BIC | 20.0 | -1.4 | 7.3 |
|  | 60.0 | 7.0 | 4.2 |
|  | 800 | 3.1 | 4.5 |
|  | 16000 | -2.8 | 4.1 |
| DOR | 3.00 | -2.8 | 11.4 |
|  | 9.00 | 2.9 | 7.4 |
|  | 120 | 2.5 | 5.6 |
|  | 2400 | 0.8 | 3.0 |
| RAL | 10.0 | 3.6 | 5.3 |
|  | 30.0 | 8.5 | 6.0 |
|  | 400 | 6.9 | 6.1 |
|  | 8000 | -0.5 | 4.2 |

### 3.4. Carryover/Crosstalk

No carryover was observed for any analyte or internal standard following the injection of a blank sample immediately following the highest calibration standard. Cross talk was also evaluated by extracting a high standard without internal standard being added. No peak was seen in the internal standard channel showing no positive bias in the internal standard response at higher analyte concentrations.

### 3.5. Recovery/Matrix Effects

Three experiments were performed to evaluate recovery and matrix effects. Peak area responses of the analytes and the internal standard extracted from human plasma at the low, mid, and high QC concentrations (pre-extracted) were compared with blank extracts spiked with analyte and internal standard corresponding to 100% recovered concentrations (post-extracted) and neat solution samples (un-extracted) corresponding to 100% recovered concentrations. True recovery was calculated as the ratio of the mean peak area response of the compound in pre-extracted QCs to the mean peak area response in post-extracted QCs. The range of recovery associated with this assay was as follows: 99.1%-110% (BIC), 91.5%-111% (DOR), and 97.0%-111% (RAL) for the three QC concentrations evaluated. Recovery of internal standards was 99.4%-101% (BIC-d_5_), 97.5%-104% (^13^C_6_-DOR), and 94.0%-104% (RAL-d_6_). The recovery and matrix effects were consistent between the analytes and internal standard throughout the concentration range evaluated.

An additional evaluation of matrix effects was performed using the experiments previously described by Matuszewski [6]. Six different lots of blank plasma were each spiked at the low, mid, and high QC levels and extracted in triplicate with the average peak area ratios from the analyses being plotted against nominal QC concentrations for each of the six lots evaluated. The slope values from these six curves were compared. The %CV from the six slope values for all three analytes were <5% indicating the lack of matrix effect related to different lots of plasma used in the extraction. These data are presented in Table 2.

**Table 2:** BIC, DOR, and RAL Slope Comparison and LLOQ QC Precision and Accuracy from Human Plasma Samples Prepared in 6 Different Lots

| Analyte | Parameter | Lot 1 | Lot 2 | Lot 3 | Lot 4 | Lot 5 | Lot 6 | %CV (Slope) |
| --- | --- | --- | --- | --- | --- | --- | --- | --- |
| BIC | Slope | 0.00096 | 0.00088 | 0.00089 | 0.00088 | 0.00087 | 0.00089 | 3.3 |
|  | LLOQ %CV | 3.3 | 3.7 | 7.1 | 4.8 | 1.3 | 9.5 |  |
|  | LLOQ %Bias | -3.3 | -7.2 | -2.0 | -7.5 | -3.3 | -6.8 |  |
| DOR | Slope | 0.00292 | 0.00274 | 0.00270 | 0.00252 | 0.00264 | 0.00263 | 4.8 |
|  | LLOQ %CV | 4.1 | 9.8 | 1.6 | 3.0 | 3.3 | 8.0 |  |
|  | LLOQ %Bias | 7.9 | 0.6 | -3.8 | 1.6 | -5.4 | 7.4 |  |
| RAL | Slope | 0.00128 | 0.00121 | 0.00121 | 0.00123 | 0.00126 | 0.00128 | 2.4 |
|  | LLOQ %CV | 6.3 | 2.6 | 2.8 | 7.1 | 7.6 | 6.3 |  |
|  | LLOQ %Bias | 6.2 | 16.0 | 8.0 | 6.4 | 5.4 | 4.0 |  |

Blank plasma from the six selectivity lots were spiked at the LLOQ QC concentration (n=3) and quantified against the calibration curve to determine if assay performance was consistent in different lots of plasma. Precision (%CV) and accuracy (%Bias) were within the ±20% acceptance criteria, showing that the use of different lots of plasma do not impact accurate quantitation of BIC, DOR, and RAL. These data are presented in Table 2.

### 3.6. Dilutions

A dilution QC (60,000 ng/mL (BIC), 9,000 ng/mL (DOR), and 30,000 ng/mL (RAL)) was evaluated as part of the method validation. The dilution QC was diluted with blank human plasma 10x prior to extraction. All three analytes met 15% precision and accuracy requirements demonstrating the ability to dilute samples with concentrations up to 60,000 ng/mL (BIC), 9,000 ng/mL (DOR), and 30,000 ng/mL (RAL) into the validated calibration range.

### 3.7. Stability

#### 3.7.1 Extract/Reinjection Reproducibility

Plasma extracts from QC samples at the low (60.0 ng/mL (BIC), 9.00 ng/mL (DOR), 30.0 ng/mL (RAL)) and high (16,000 ng/mL (BIC), 2,400 ng/mL (DOR), 8,000 ng/mL (RAL)) concentrations from a prior validation run were stored in the autosampler at 15⁰C for 3 days. To test extract stability, after 3 days, these low and high QC (n=3) were reinjected and quantified against the original calibration curve. The resulting concentrations back to the nominal levels for the low and high QC respectively were 6.8% and −4.0% (BIC), 7.8% and −2.5% (DOR), and 10.3% and −3.1% (RAL) deviation. For reinjection reproducibility, the same extracts at the low and high QC concentrations (n=3), along with the duplicate calibration curves, stored for 3 days at 15°C were reinjected and analyzed. Comparisons of these concentrations to nominal concentrations for the low and high QC resulted in 11.3% and −3.1% (BIC), 8.1% and −6.8% (DOR), and 10.9% and −2.0% (RAL), respectively. Both stability tests met the acceptance criteria of ±15%.

#### 3.7.2 Stock/Working Solution Stability

Long-term stability of stock solutions was tested by comparing previous prepared solutions that were stored at −80°C to solutions that were freshly prepared. The percent difference from the stored stock was 2.3% (BIC), 0.1% (DOR), and −2.4% (RAL) showing that BIC and DOR are stable for up to 158 days (storage at the time of investigation) and up to 591 days for RAL when stored at −80°. The stock stability of BIC and DOR is ongoing. Stability of stock solutions was also evaluated at room temperature. Following initial stock solution preparation, an aliquot of each of the stock solutions was transferred into a tube and stored at room temperature for either 24 hours (BIC, DOR) or 18 hours (RAL), while a second aliquot was transferred to a separate tube and placed back at −80°C. After 24 or 18 hours the frozen aliquot was removed from the freezer and analyzed. The % difference was 5.6% (BIC), 6.8% (DOR), and 0.30% (RAL) demonstrating that all stock solutions are stable at these conditions.

In addition, we evaluated the stability of the spiking solutions prepared in 50:50 methanol:water. Fresh spiking solutions prepared at the low and high QC concentrations were compared to solutions that were stored at −80°C for 155 days. The percent differences between these solutions were 0.1% (BIC), 3.1% (DOR), and 0.6% (RAL) for the low QC concentration and 0.2% (BIC), −5.7% (DOR), and −3.5% (RAL) for the high QC concentration demonstrating stability when stored at −80°C.

#### 3.7.3 Freeze-Thaw/Room-Temperature Stability

The stability of BIC, DOR, and RAL in human plasma was evaluated under various storage conditions. Human plasma QC samples at the low (60.0 ng/mL (BIC), 9.00 ng/mL (DOR), 30.0 ng/mL (RAL)) and high (16,000 ng/mL (BIC), 2,400 ng/mL (DOR), 8,000 ng/mL (RAL)) QC concentrations were allowed to sit at room temperature for approximately 21 hours (BIC, DOR) and 24 hours (RAL). These QCs were assayed in triplicate and compared to nominal concentrations. The room temperature exposed QCs showed 1.0% and −4.0% (BIC), 1.0% and 0.4% (DOR), and 11.3% and −3.3% (RAL) deviation from nominal at the low and high QC concentrations, respectively, meeting the acceptance criteria of ±15%. The stability of BIC, DOR, and RAL in human plasma after 3 freeze/thaw cycles between room temperature and −80°C was evaluated. The first cycle was frozen for at least 24 hours and all other cycles were frozen for at least 12 hours between each cycle. Low and high QC samples were assayed in triplicate after completing 3 freeze/thaw cycles. The freeze/thaw QCs showed a 7.8% and −0.4% (BIC), 2.4% and 0.8% (DOR), and 14.0% and −0.2% (RAL) deviation from nominal at the low and high QC levels, respectively, meeting the acceptance criteria of ±15%.

#### 3.7.4 Storage Stability

Long-term stability in human plasma for BIC, DOR, RAL was performed using previously prepared low QC and high QC samples that were placed in −80°C for 154 days. Additional longer-term assessments will be performed to establish longer matrix stability. The concentrations of the stored QC samples were within ±6.1% (BIC), ±5.2% (DOR), and ±6.8% (RAL) of the theoretical concentrations, which demonstrated stability by being within the ±15% acceptance criteria.

#### 3.7.5 Whole Blood Stability

Low (60.0 ng/mL (BIC), 9.00 ng/mL (DOR), 30.0 ng/mL (RAL)) and high (16,000 ng/mL (BIC), 2,400 ng/mL (DOR), 8,000 ng/mL (RAL)) QC samples were prepared in whole blood. Multiple aliquots of each QC were transferred into individual tubes. An aliquot of each QC level was immediately centrifuged and the plasma layer was removed (T=0 sample). This processing of whole blood into plasma was repeated following storage at room temperature and in an ice bath for 1 and 4 hours The resulting plasma samples were extracted and the peak area ratios compared to the T=0 sample. BIC and DOR were stable for up to 4 hours for both room-temperature and on ice conditions, with % difference for room temperature [low, high QC] from the T=0 as follows: −4.9, −6.2% (BIC) and 6.8%, 0.1% (DOR) and %difference for on ice [low, high QC] as: −6.2, −10.0% (BIC) and −13.0%, 5.6% (DOR). RAL was stable for up to 1 hour at both room temperature and on ice with the % difference for room temperature [low, high QC] from the T=0 as follows: 4.3%, −3.6% and the % difference for on ice [low, high QC] as: −1.2, −4.4%. The low concentration stability was outside of the limit of acceptability for both room temperature and the ice bath with the % difference from T=0 being 15.6% in these conditions The high QC was stable for up to 4 hours with the %difference [room temperature, ice] from T=0 being 5.9% and 5.0%.

## 4. Conclusions

An LC-MS/MS assay has been developed for the analysis of bictegravir, doravirine, and raltegravir in human plasma. This method was fully validated according to the FDA guidance for industry standards in human plasma demonstrating linearity and sensitivity over each analyte’s calibration range along with acceptable accuracy and precision. This assay increases sensitivity to aid in monitoring adherence for the quantification of BIC, DOR, and RAL in human plasma. Our assay’s sensitivity is 100-130-fold lower than typical trough concentrations (Ctrough) observed in HIV infected patients at the end of the FDA approved dosing interval for BIC and DOR [7, 8]. For RAL, sensitivity is 3.5- and 7-fold lower than the typical Ctrough for once and twice daily dosing, respectively [9, 10]. Given our assay’s sensitivity, the application of this method to support clinical pharmacokinetic investigations or adherence and therapeutic drug monitoring for these 3 antiretrovirals is appropriate.

## Acknowledgements

This research was supported in part by the University of North Carolina at Chapel Hill Center for AIDS Research (CFAR), an NIH funded program P30 AI50410.

